# Fluorescent Sp^3^ Defect-Tailored Carbon Nanotubes Enable NIR-II Single Particle Imaging in Live Brain Slices at Ultra-Low Excitation Doses

**DOI:** 10.1101/636860

**Authors:** Amit Kumar Mandal, Xiaojian Wu, Joana S. Ferreira, Mijin Kim, Lyndsey R. Powell, Hyejin Kwon, Laurent Groc, YuHuang Wang, Laurent Cognet

## Abstract

Cellular and tissue imaging in the second near-infrared window (NIR-II, ∼1000 - 1350 nm) is advantageous for *in vivo* studies because of low light extinction by biological constituents at these wavelengths. However, deep tissue imaging at the single molecule sensitivity has not been achieved in the NIR-II window due to lack of suitable bio-probes. Single-walled carbon nanotubes have emerged as promising near-infrared luminescent molecular bio-probes; yet, their inefficient photoluminescence (quantum yield ∼1%) drives requirements for sizeable excitation doses (∼1-10 kW/cm^2^) that are significantly blue-shifted from the NIR-II region (<850 nm) and may thus ultimately compromise live tissue. Here, we show that single nanotube imaging can be achieved in live brain tissue using ultralow excitation doses (∼100 W/cm^2^), an order of magnitude lower than those currently used. To accomplish this, we synthesized fluorescent sp^3^-defect tailored (6,5) carbon nanotubes which, when excited at their first order excitonic transition fluoresce brightly at ∼1160 nm. The biocompatibility of these functionalized nanotubes, which are wrapped by state-of-the-art encapsulation agents (phospholipid-polyethylene glycol), is demonstrated using standard cytotoxicity assays. Single molecule photophysical studies of these biocompatible nanotubes allowed us to identify the optimal luminescence properties in the context of biological imaging.

## Introduction

Optical imaging of live biological tissue is a promising and actively developing field with major expected impacts on the fundamental understanding of various biological processes but also on diagnostic, therapeutic and health care. Compared to other imaging modalities, optical imaging offers the combination of exquisite properties including sensitivity (down to single molecule level) and resolution (molecular) while being compatible with living sample analysis. One major drawback however is the limited penetration of optical photons into biological tissue. To date, most high-resolution and/or high-sensitivity imaging methods have been developed at visible wavelengths (i.e. ∼400-700 nm) where light is strongly scattered by the biological constituents. ^[1]^ Strong efforts have also been made to reach the short near-infrared (NIR-I) wavelength range (∼700-1000 nm), also called the diagnostic or therapeutic window, where light better propagates than visible light into biological tissues. This led to the development of a variety of non-invasive optical imaging of techniques reaching the single molecule level in living tissue, or, with poorer sensitivities in the small animal. More recently, two additional red-shifted optical windows have been identified and referred as the NIR-II (∼1000-1350 nm) and NIR-III (∼1550-1900 nm) optical windows with distinct advantages in terms of light scattering and absorption depending on photon energy. ^[2,3]^ In particular, the NIR-II window is attracting high attention for the combination of low light scattering and absorption by the tissues, resulting in good light penetration depth.

In order to fully exploit the high resolution - high sensitivity properties of optical imaging, one generally introduces optical nanoprobes into the tissue that are small to access complex biological environments and display intense optical signals, chiefly fluorescence signals. Unfortunately, nanoparticles that are optically active into the NIR-II region are uncommon, ^[4-7]^ especially for single particle applications, which require bright emitters. Yet, this would enable obtaining invaluable information about detailed biological processes in complex tissues with nanoscale resolutions. Soon after the discovery of their photoluminescence (PL) in the NIR,^[8]^ single-walled carbon nanotubes (SWCNTs) were identified as unique biological probes^[9]^ due to their unparalleled PL stability and brightness in the NIR and latter complemented by their unique 1D morphology which allow efficient access and dissemination in tissues.^[10,11]^ At the ensemble level, SWCNTs can be imaged in live cells, tissue and even animals over long periods of time in the NIR-I optical window but also in the NIR-II and even in the NIR-III optical windows.^[9,12]^ At the single molecule level, the task is intrinsically difficult due to the relatively weak SWCNT signals which must compete with any faint source of tissue autofluorescence, light scattering and absorption. In addition, light excitation necessary to achieve efficient single SWCNT PL must be compatible with live tissue integrities. ^[13]^ Due to these constraints, imaging low concentrations of SWCNTs, ^[14,15]^ notably single SWCNTs ^[10]^ into live tissue remains challenging. Note that reduced sensitivity and higher noise of detectors at NIR wavelengths complicates the task. In this context, a promising approach would be to enhance simultaneously SWCNT emission and absorption properties to achieve high signal-to-noise ratio imaging of single SWCNTs in the NIR-II window using minimal excitation intensities.

SWCNTs have low luminescence quantum yield (QY) ^[8]^ due to non-radiative mechanisms and photoluminescence quenching by structural defects and environmental quencher. ^[16-20]^ Mobile excitons, which are the energy carriers in a SWCNT, can diffuse over one to several hundreds of nanometers along the nanotube, ^[17,21]^ and thus is affected by any PL quenching defect along the SWCNT length or at its ends. In order to improve this low luminescence QY, an elegant approach consists in covalently attaching low density chemical functional groups to the SWCNTs, e.g., alkyl and aryl groups, leading to the creation of sp^3^ defects that strongly localize the band-edge excitons into quantum states (E_11_^−^) located 100-300 meV below the first bright band-edge exciton level (E_11_).^[22-28]^ These sp^3^ quantum defects, instead of quenching the SWCNT PL photoluminescence, channel the excitons to the emissive defects leading to more than an order-of-magnitude brightening effect at the ensemble level.^[24]^ Even ultrashort, essentially non-emissive SWCNTs become brightly fluorescent with this method.^[29]^ Localized emission at individual defects sites is directly observed using wide-field PL imaging and even with super-resolution precisions.^[29]^ Interestingly, the E_11_^−^ emission of functionalized SWCNTs being red-shifted from the strongest excitonic transition (E_11_), one can take advantage of this possibility to generate E_11_^−^ emission by direct excitation at the E_11_ transition.^[23]^

Here, we show that bright, biocompatible *p*-nitroaryl functionalized SWCNTs (f-SWCNTs) encapsulated in phospholipid-polyethylene glycol (PL-PEG) are suitable for application in single molecule bio-imaging. We demonstrate imaging of these f-SWCNTs in live organotypic brain tissues at the single nanotube level and compare the results with the biocompatible unfunctionalized analogue (unf-SWCNTs). The f-SWCNTs enable high signal-to-noise ratio imaging within the NIR-II region (985 nm excitation, 1160 nm emission) using excitation intensities that are one order of magnitude less than those required for the biocompatible unf-SWCNTs (985 nm excitation and emission).

## Materials and Methods

### unf-SWCNT encapsulation in phospholipid-polyethylene glycol (PL-PEG)

CoMoCAT SG65i nanotubes were suspended by 0.5 % w/v PL-PEG (MPEG-DSPE-5000, Layson Bio. Inc., Arab, AL, USA) in deuterium oxide (D_2_O). 1 mg of raw SWCNTs was added to 10 mL of the PL-PEG solution. The mixture was homogenized and sonicated with a tip sonicator (Misonix-XL 2000, 20 W output) for 8 min in an ice bath. Nanotube bundles and impurities were precipitated by centrifugation (Beckman Coulter, 50.2 Ti rotor) at 6000 rpm for 60 min at room temperature. The supernatant was collected and stored at 4°C until further use. The absorption spectrum (400-1000 nm) was recorded for every suspension with a spectrometer (Cary 5000). The concentration of the PL-PEG-coated SWCNT was estimated to be 0.36 μg mL^−1^ based on the optical absorbance at 808 nm.^[30]^

### Covalent incorporation of fluorescent sp^3^ defects in SWCNTs

SWCNTs were covalently functionalized by *p*-nitro aryl groups (-C_6_H_5_NO_2_) using a diazonium reaction in oleum. 3 mg of raw CoMoCAT SG65i SWCNTs (99%, Lot No. SG65i-L39, SouthWest NanoTechnologies) were added to 10 mL of oleum (20% free SO_3_, Sigma-Aldrich) and stirred overnight. (*Safety note: Oleum is extremely acidic and corrosive. The reaction must be performed within a fume hood with protective gloves and masks*.) 0.86 mg of sodium nitrite (NaNO_2_; analytical standard, Sigma-Aldrich) and 1.75 mg of *p*-nitroaniline (analytical standard, Sigma-Aldrich) were added to the SWCNT/oleum suspension at a molar ratio of SWNCT carbon relative to *p*-nitroaniline of 20:1. The mixture was heated to 75°C using an oil bath, and 2.5 mg of azobisisobutyronitrile (98%, Sigma-Aldrich) was then added to initiate the reaction. After 20 min of stirring at 75°C, the system was removed from the oil bath and cooled to room temperature. The SWCNT-oleum suspension was added dropwise to 250 mL of a 2.0 M aqueous solution of sodium hydroxide (97%, Sigma-Aldrich) until the solution reached pH 8. (*Safety note: This neutralization step must be performed slowly with stirring within a fume hood in order to avoid the production of excessive heat or toxic fumes.)* The neutralized SWCNT suspension was filtered by vacuum over a 80 nm pore size track-etched polycarbonate membrane and the filter cake was rinsed with copious water and ethanol to remove unreacted chemicals. The f-SWCNTs obtained on the filter paper were dried in vacuum for 1 h and then re-dispersed in 1% w/v DOC-D_2_O by probe sonication with a 3.2 mm diameter microtip at 8 W for 1 h.

### Direct and exchanged f-SWCNT

The f-SWCNTs were either dispersed directly in 0.5% w/v PL-PEG by probe sonication for 1 h at 4 W, or first dispersed in 1% w/v DOC-D_2_O for 1 h, again by sonication at 4 W, then exchanged with 0.5% PL-PEG-D_2_O by pressure filtration (Amicon Stirred Cell, EMD Millipore). The photoluminescence of nanotube solutions was collected using a NanoLog spectrofluorometer (HORIBA Jobin Yvon). The samples were excited with a 450 W Xenon source dispersed by a double-grating monochromator. The PL spectra were then collected using a liquid-N_2_ cooled linear InGaAs array detector. The slit widths for both the excitation and emission were set at 10 nm.

### Cell culture studies

Hela cells were cultured on microscope coverslips in Dulbecco’s Modified Eagle Medium (DMEM) supplemented with streptomycin (100 g/mL), penicillin (100 U/mL), and 10% bovine serum in a 95% humidified atmosphere, 5% CO_2_, at 37°C. Cells were cultured every three to four days and used up to passage 20.

### Cell proliferation assay

Hela cells were incubated with SWCNTs at a final concentration of 5 μg/mL in DMEM for one day or four days at 5% CO_2_ at 37°C. The cells were then cleaved from the coverslips using trypsin (1%) and harvested by centrifugation at 1200 rpm for 5 min, after which the cells were suspended in 1 mL of DMEM. For cell counting, using microscopy at 10X magnification, a 10 μL drop of the sample was placed at the center of hemocytometer and covered with a coverslip. The sample was mounted on a homemade microscope with white light illumination at room temperature and cells were counted. The cell number was calculated and normalized by the control cell number without nanotube incubation.

### Cell viability assay

For the cell viability assay, two sets of Hela cells were grown in a 96-well tissue culture plates for either one or four days. In each well, 200 μL of the cell sample was incubated in 20 μL of Cell Proliferation Reagent WST-1 (Sigma Aldrich) for 1.5 h at room temperature to lyse the cells. After the incubation period, the resulting formazan dye was quantitated with a scanning multi-well plate spectrophotometer (Biorad iMark plate reader). The absorbance directly correlates to the number of viable cells. Blank absorption collected only from the same volume of culture media, and WST-1 reagent (in the absence of cells) was subtracted from each absorbance value. Control cells were prepared by following the same protocol, but without SWCNTs administration. The viable cell fractions (percentages) were calculated by normalizing the cell number with nanotube incubation by the control cell number without nanotube incubation.

### Single Nanotube Photoluminescence Imaging

SWCNT PL imaging was performed with an inverted microscope equipped with a 60X (1.40 NA) objective and two detection arms separated by a 50/50 beam splitter. A tunable Ti-sapphire laser (Spectra Physics, Model 3900S) was to excite the (6,5) SWCNTs at their K-momentum exciton-phonon sideband (845 nm). In addition, a 985 nm diode laser was used to resonantly excite the functionalized (6,5)-SWCNTs at their first order excitonic transition (E_11_). Fluorescence was collected with the same objective and imaged on an InGaAs camera (Ninox, Raptor Photonics), in order to produce wide-field images of individual SWCNTs. A dichroic mirror (FF875-Di01, Semrock, Rochester, NY, USA) and the combination of long- and short-pass emission filters (ET900LP, Chroma Technology Corp., Bellows Falls, VT, USA; FESH1000, Thorlabs SAS, Maisons-Laffitte, France) were used to illuminate the sample and detect SWCNT fluorescence at 985 nm. Images of SWCNTs were recorded with 30 ms integration time per frame. A dichroic mirror (FF875-Di01, Semrock) in combination with long-pass emission filters (RazorEdge 1064, Semrock) for (6,5) f-SWCNTs was used to illuminate the sample and select PL from the E_11_^−^ state only. PL from E_11_ and the E_11_^−^ were detected with the InGaAs camera. For imaging immobilized SWCNTs, the suspended SWCNT samples were spin-coated on a microscope glass coverslip pre-coated with polyvinylpyrrolidone. In this work, unless otherwise stated, the PL of an individual SWCNT is defined as the signal integrated over the pixels from the nanotube image. In Figure 3A, the reported luminescence brightness of the unf-SWCNTs was multiplied by 1.034 to account for the difference of detection efficiency of the InGaAs camera between 985 nm and 1160 nm.

**Figure 1:**
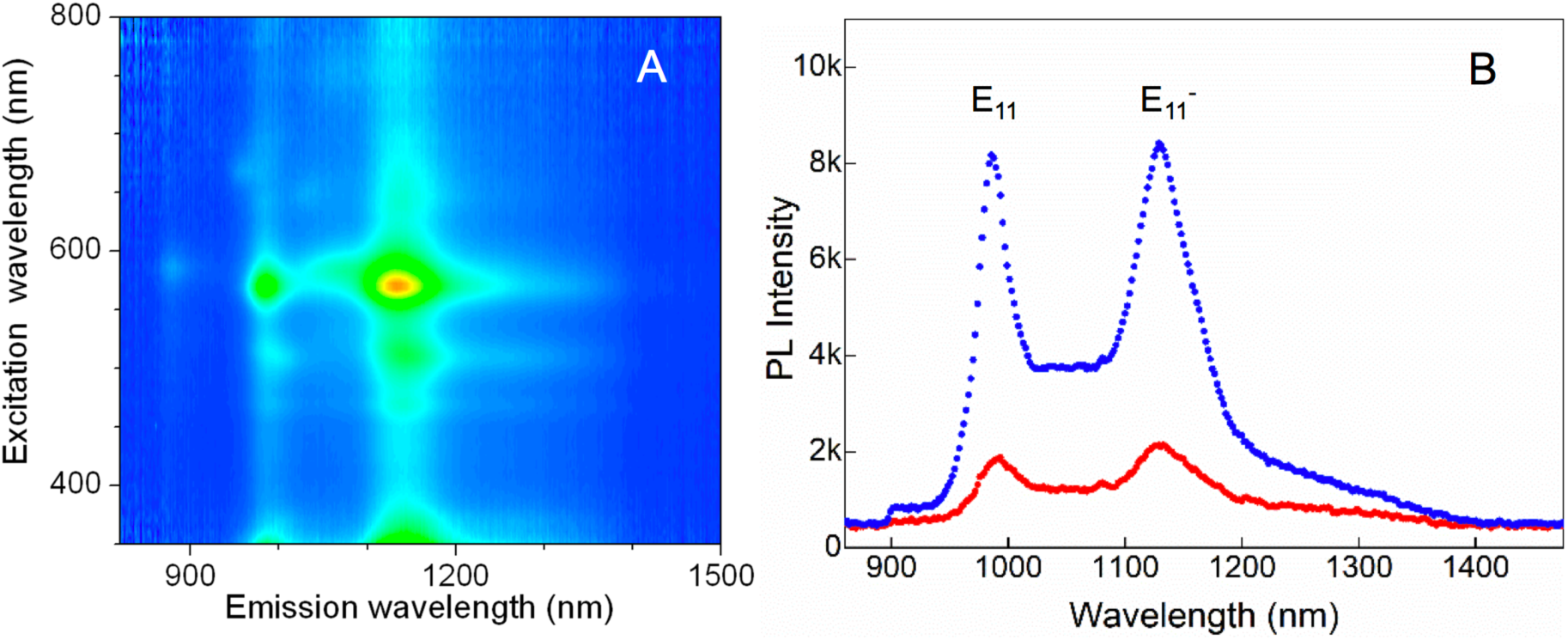
(A) Excitation-emission photoluminescence map of p-nitroaryl tailored (6,5)-SWCNTs with native PL (E_11_) and defect emission (E_11_^−^) peaks at 985 nm and 1160 nm, respectively. (B) Emission spectra of the f-SWCNTs prepared by two different methods: (red) directly stabilized in PL-PEG and (blue) stabilized in DOC then exchanged into PL-PEG by pressure filtration. The nanotube concentrations were adjusted to 0.36 μg mL^−1^. The excitation wavelength was 565 nm.

**Figure 2:**
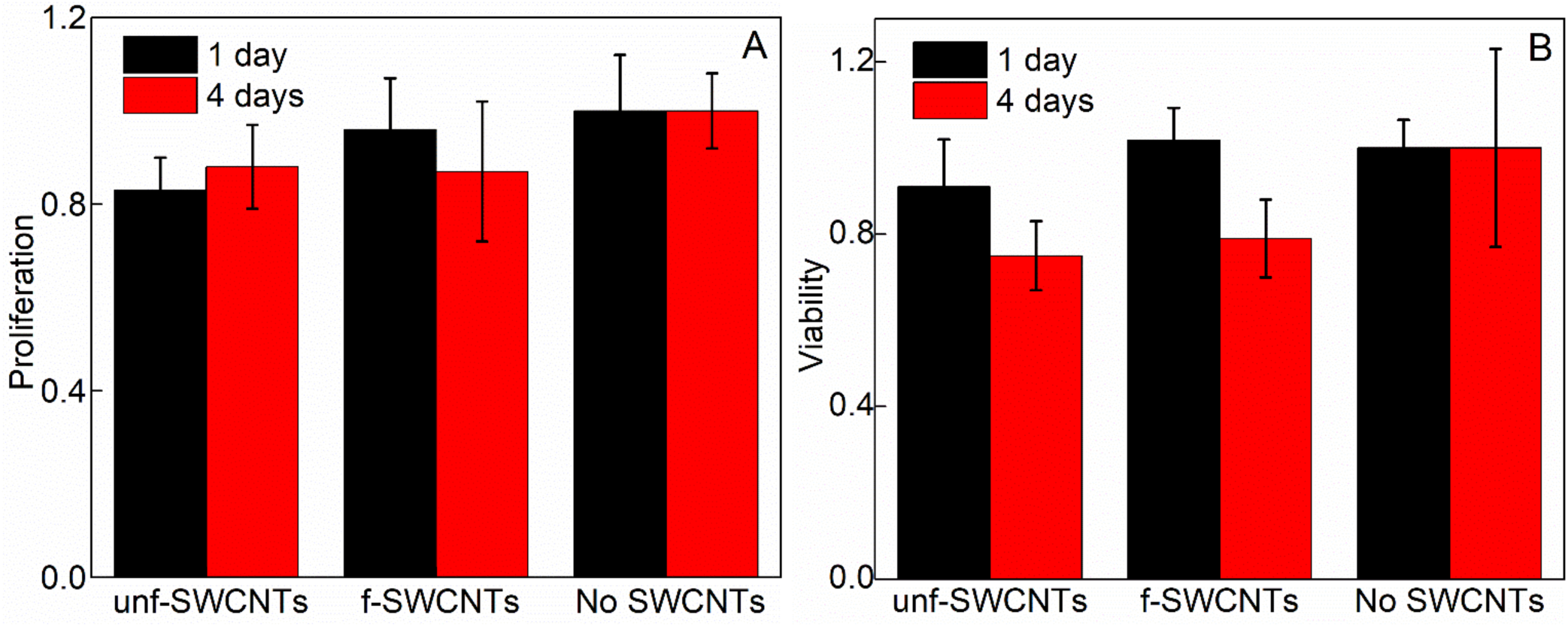
Live cell biocompatibility test of unf- and f-SWCNTs. (A) Cell proliferation investigation. SWCNT concentration in the cell culture: 5 μg/mL. (B) Cell viability study with WST-1. SWCNT concentration in the cell culture: 20 μg/mL. In both assays, cell proliferation and viability were compared with the blank control sample without SWCNTs.

**Figure 3:**
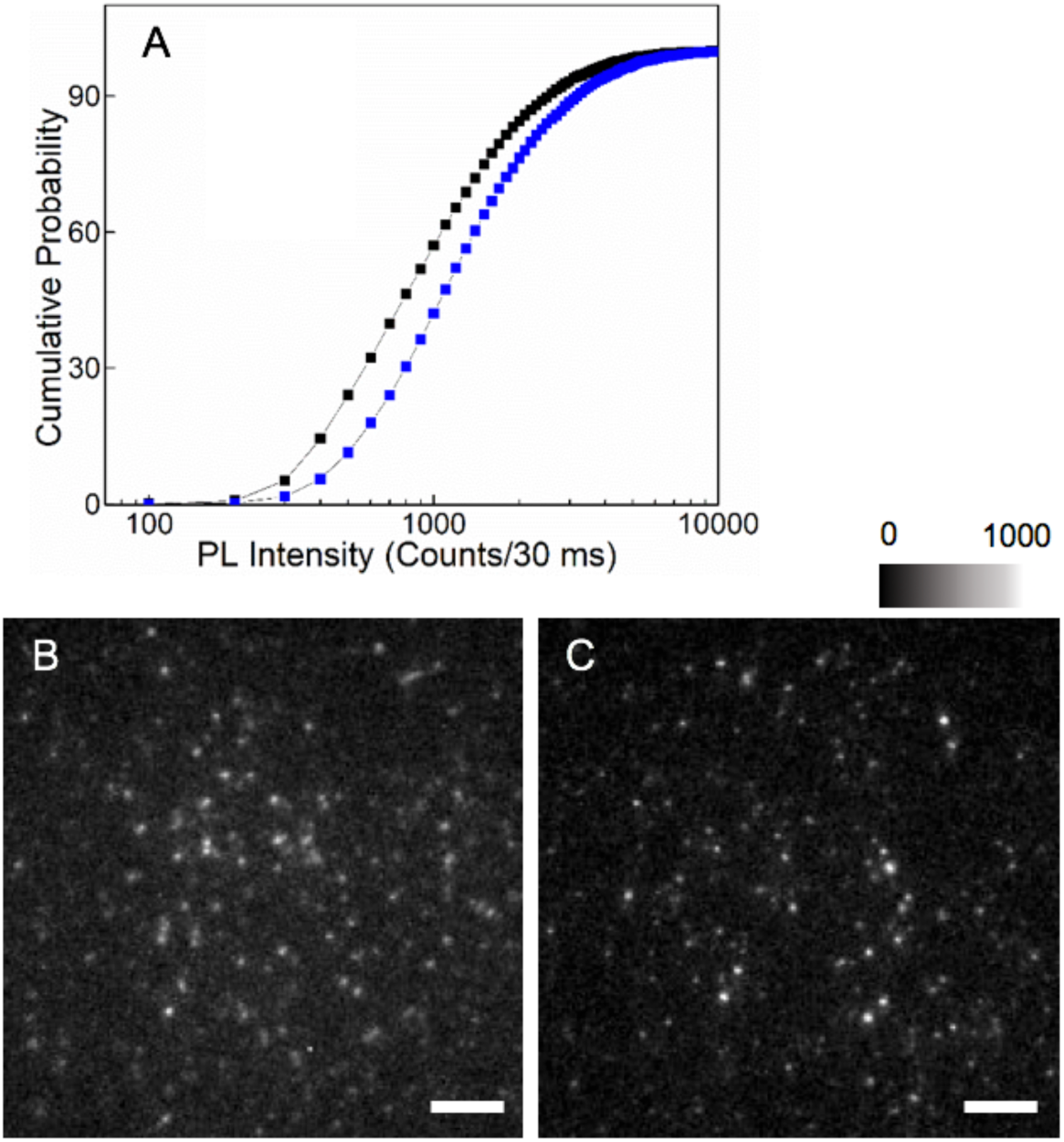
(A) Cumulative distribution of the photoluminescence of unf-SWCNT (E_11_, black line) and f-SWCNT (E_11_^−^, blue line) resulting from 568 nm excitation at 0.3 kW/cm^2^. PL intensity corresponds to integrated signals from individual SWCNT images. (B) Image of E_11_ photoluminescence emission from individual unf-SWCNTs recorded using a 900 nm long pass filter (C) Image of E_11_^−^ photoluminescence from individual f-SWCNTs recorded using a 1064 nm long pass filter. The scale bar is 10 μm in B and C.

### Organotypic Tissue Preparation

Organotypic slice cultures were prepared as previously described.^31^ Briefly, 350-μm hippocampal slices were obtained from postnatal day 5 (P5) to P7 Sprague-Dawley rats using a McIlwain tissue chopper, and were placed in a pre-heated (37°C) dissection medium containing 175 mM sucrose, 25 mM D-glucose, 50 mM NaCl, 0.5 mM CaCl_2_, 2.5 mM KCl, 0.66 mM KH_2_PO_4_, 2 mM MgCl_2_, 0.28 mM MgSO_4_-7H_2_O, 0.85 mM Na_2_HPO_4_-12H_2_O, 2.7 mM NaHCO_3_ and 0.4 mM HEPES, 2 × 10-5% phenol red, pH 7.3 (all products from Sigma Aldrich unless specified). After 25 min of incubation, slices were transferred onto white hydrophilic polytetrafluoroethylene membranes (0.45 μm; Millipore FHLC) set on Millicell Cell Culture Inserts (Millipore, 0.4 mm; 30 mm diameter), and cultured for up to 14 days on multiwell-plates at 35°C/5% CO_2_ in a culture medium composed of 50% Basal Medium Eagle, 25% Hank’s balanced salt solution 1X (with MgCl_2_/with CaCl_2_), 25% heat-inactivated horse serum, 0.45% D-glucose and 1 mM L-glutamine (all products from Gibco unless specified). The medium was changed every 2-3 days.

### Organotypic Tissue Imaging

The tissue slices were first incubated with SWCNT for ∼2 h, and then the slices were mounted in a ludin chamber for image acquisition. The ludin chamber was filled with artificial cerebrospinal fluid solution (126 mM NaCl, 3.5 mM KCl, 2 mM CaCl_2_, 1.3 mM MgCl_2_, 1.2 mM NaH_2_PO_4_, 25 mM NaHCO_3_ and 12.1 mM glucose; gassed with 95% O_2_: 5% CO_2_; pH 7.35) at room temperature. (6,5)-SWCNTs were selectively excited within the tissue at 845 nm by a tunable Ti-Sa laser (Spectra Physics, Model 3900S) and 985 nm by a diode laser (Thorlabs). The incident laser beam, circularly-polarized at the sample to ensure that SWCNTs are excited regardless of their orientation in the sample plane, was focused into the back aperture of an objective (Nikon, 60x, NA 1.0) mounted on an upright microscope (Nikon TiE eclipse; Tokyo, Japan). The emission was collected with the same objective and imaged on the same camera to produce wide-field images of individual SWCNTs within the tissue. A dichroic mirror (FF875-Di01; Semrock, Rochester, NY, USA) and the combination of long-pass emission filters (ET900LP, Chroma Technology Corp. and RazorEdge 1064, Semrock) were used to illuminate and detect PL of SWCNTs inside deep tissues. Images of SWCNTs were recorded with 30 ms integration time per frame.

## Results and Discussion

### Biocompatible functionalized nanotubes (the f-SWCNT)

The f-SWCNTs were prepared by covalent functionalization of CoMoCAT SG65i nanotubes with *p*-nitroaryl groups using aryldiazonium salts in oleum. The functionalized nanotubes were collected by vacuum filtration and re-dispersed in 0.5% w/v PL-PEG or 1% w/v DOC/D_2_O by probe sonication (see Materials and Methods). Successful incorporation of the fluorescent sp^3^ defects in (6,5)-SWCNTs is directly confirmed from the observation of the intense defect PL (E_11_^−^) at 1160 nm, which is redshifted from the the native PL (E_11_) from unfunctionalized segments of the (6,5) nanotubes.

We then suspended the f-SWCNT in PL-PEG, which improves the biocompatibility of SWCNTs. ^[32,33]^ Note that while the use of surfactants such as sodium deoxycholate or sodium dodecylbenzene sulfonate result in solutions with brighter nanotubes, ^[32]^ these surfactants are not applicable to in-vivo imaging, as they are detrimental to cell membrane integrity. Two different methods were explored for SWCNT dispersion from the filter cake. The first one was to directly disperse the SWCNTs in PL-PEG while the second one consisted in first dispersing the f-SWCNTs in 1% DOC-D_2_O then exchanging the surfactant with 0.5% PL-PEG-D_2_O by pressure filtration. Interestingly, the second method led to samples having higher PL brightness than the first one when comparing identically concentrated nanotube solutions (Figure 1B). This is probably due to the superior efficiency of DOC to individualize small diameter nanotubes from dry into aqueous environments.^[34]^ Owing to their biocompatibility and photoluminescence brightness, we exclusively chose the exchanged f-SWCNT for all following experiments. To monitor their suitability and usefulness for biological applications, we next performed biocompatibility and cytotoxicity studies toward live cells using the standard Hela cell line.

### Cytotoxicity Experiments

In order to address the potential cytotoxicity of f-SWCNTs, we first performed a cell proliferation study upon incubation with a final nanotube concentration of 5 μg/mL. The cell numbers were counted after 1 day and after 4 days using a hemocytometer under a white light microscope. Two controls were also used: cell incubation with unf-SWCNTs encapsulated in PL-PEG (using the same concentration) and without nanotube application. In these experiments, the initial quantity of Hela cells was adjusted to 8×10^4^ cells/mL. In accordance with previous studies, unf-SWCNTs encapsulated in PL-PEG showed minimal cell proliferation inhibition at both time points as compared to control cells (more than 80%). ^[32]^ Importantly, f-SWCNTs showed an identical behavior (Figure 2A).

In order to check cell viability, we complemented this analysis by the standard WST-1 assay that is used widely for analysis of acute cytotoxicity.^[35]^ In this case, HeLa cells were incubated (5×10^4^ cells/mL) with unf- and f-SWCNTs, with a final concentration of 20 μg/mL. After 1 day and after 4 days, viable cells were identified using WST-1 staining and cell viability was calculated (see Materials and Methods). Cell viabilities were greater than 90% after 1 day and 70% after 4 days of incubation (Figure 2B). Note that in this experiment, much higher nanotube concentrations were employed than for typical *in vivo* or tissue imaging (∼0.1 to 0.2 μg/mL). We can thus conclude that the f-SWCNTs encapsulated in PL-PEG can be safely applied for imaging application in live tissue.

### Photoluminescence characterization of f-SWCNTs and comparison with unf-SWCNTs

We next compared the photoluminescence brightness of the PL-PEG wrapped unf- and f-SWCNT at the single nanotube level under different illuminations conditions. We excited both nanotubes types in solution at their second order transition (E_22_, 568 nm for (6,5) nanotubes) with identical excitation laser intensity (0.3 kW/cm^2^). In Figure 3A, the cumulative distribution of E_11_^−^ PL of individual f-SWCNTs (N=16473) is compared with that of the E_11_ PL of single unf-SWCNTs (N=22425). Under the identical excitation intensity, f-SWCNTs used in this work display 1.5-times higher photoluminescence brightness than their unfunctionalized counterpart at the single nanotube level. Note that brightening induced by sp^3^ nanotube functionalization, which was previously observed in different preparations, depends on several parameters including nanotube length, functional group nature (alkyl/aryl), starting SWCNT quality, functionalization density, etc.^[24,25,29,36,37]^ PL brightness is thus not an intrinsic parameter in sp^3^-functionalised nanotubes and should be characterized for a specific sample. In this work, brightening following functionalization is modest compared to previous reports that are better optimized for brightness, yet in the following, we will show that the f-SWCNTs offer additional benefits that are intrinsic to sp^3^-functionalisation in the context of biological imaging.

For applications in biological imaging, a visible excitation source, such as the one used in Figure 3 (568 nm) is not ideal due to limited tissue penetration depth and tissue autofluorescence which produces substantial background noise.^[3,38]^ To solve this problem in the case of unf-SWCNTs, the best option identified to date was to excite the K-momentum exciton-phonon sideband (KSB) of E_11_ which lies at 845 nm for (6,5)-SWCNTs.^[13]^ However, excitation at the KSB still resides far from the NIR-II window and it requires substantial excitation intensities (at the kW/cm^2^ level) to generate bright SWCNT PL for detection at the single molecule level. In contrast, f-SWCNTs provide a more elegant solution: direct excitation from their first order transition, E_11_ (985 nm in the case of (6,5)-SWCNTs, which is within the therapeutic window and at the edge of the NIR-II region) and detection from defect-induced E_11_^−^ (1160 nm deep in the NIR-II region). In order to evaluate the robustness of f-SWCNTs for in vivo imaging, we compared the performance of the two nanotube types as a function of excitation intensity and excitation wavelength at single-nanotube level (Figure 4). In all cases, non-linear saturation curves are observed in accordance with previous studies. Upon 845 nm excitation, both (6,5) unf-SWNCT and (6,5) f-SWCNT display rather similar photoexcitation efficiencies. Upon 985 nm excitation however, E_11_^−^ photoluminescence (at 1160 nm) for f-SWCNTs is significantly more efficient than upon 845 nm excitation. This was expected because of one order of magnitude higher absorption at the E_11_ wavelength than at the KSB. Accordingly, the saturation intensity obtained by curve fitting is ∼8 times lower upon 985 nm excitation than 845 nm (0.24 kW/cm^2^ vs 1.9 kW/cm^2^) (Figure 4A). We additionally measured the photostability of individual f-SWCNTs for up to 10 minutes under continuous excitation at 985 nm (Figure 4B). This quantitative analysis directly indicates that sp^3^-functionalized nanotubes can display similar photoluminescence rates to unfunctionalized nanotubes using significantly lower energy excitation and more importantly, with one order of magnitude less excitation intensities.

**Figure 4:**
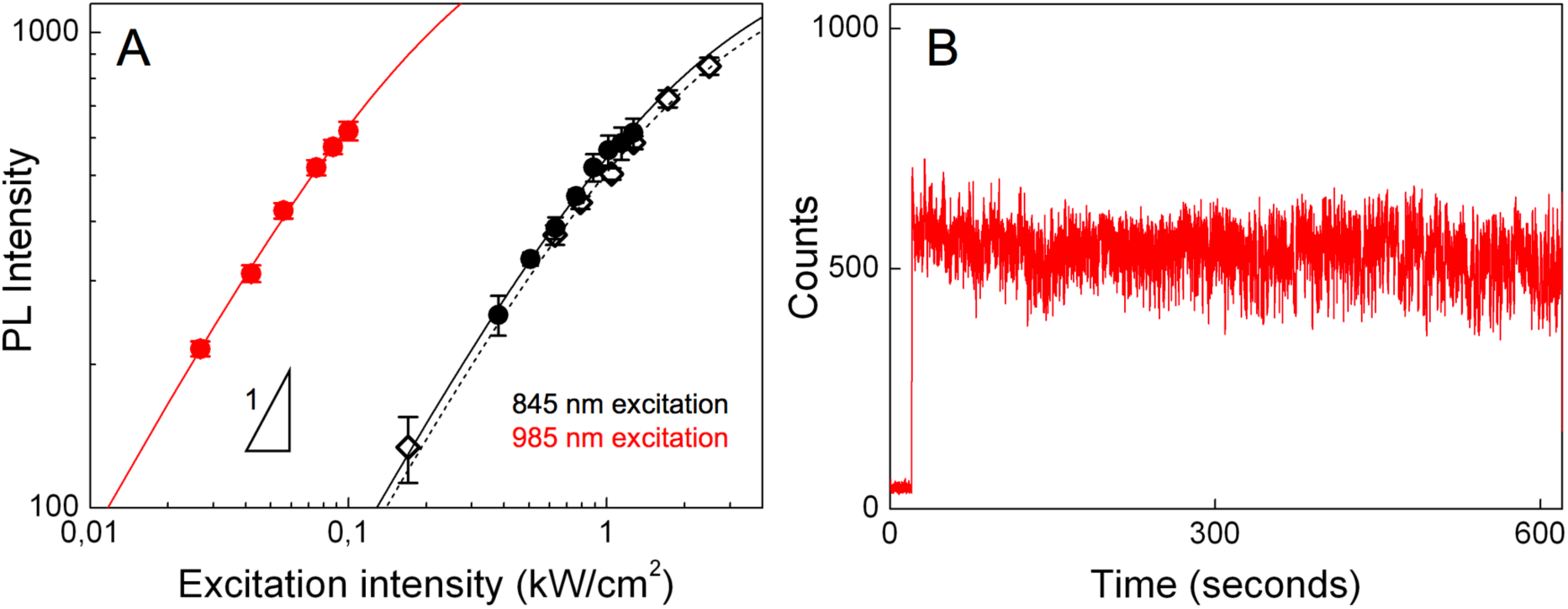
Comparison of photoluminescence brightness. (A) E_11_ photoluminescence from f-SWCNTs (black solid circles), E_11_^−^ photoluminescence from f-SWCNTs (red solid circle) and E_11_ photoluminescence from unf-SWCNTs (black open diamond) as a function of 845 nm or 985 nm laser intensity. (B) Luminescence photostability of individual f-SWCNTs for up to 10 minutes under continuous excitation at 985 nm. At ∼ 20 seconds, the excitation light is applied.

### Brain Tissue Imaging

We next used this knowledge to demonstrate that biocompatible f-SWCNTs can be instrumental for deep tissue imaging at the single nanotube level. For this, we chose live brain tissues as an archetypical system because single SWCNT detection and tracking were shown to provide unique knowledge about the brain extracellular space environment. ^[10,15,39]^ Organotypic rat brain tissue slices were prepared from P5-7 rat pups and maintained in culture medium at 35 °C/5% CO_2_ until imaging. Unf- or f-SWCNTs were delivered identically into brain tissues by incubation (120 minutes) and mounted in a ludin chamber containing the artificial cerebrospinal fluid under the microscope for imaging. Circularly polarized light was used to ensure nanotubes were excited irrespective of their orientation to the incident source. Individual nanotubes were detected in diverse areas of the brain slices, both for 845 nm excitation (unf-SWCNTs) or for 985 nm excitation (f-SWCNTs). Efficient nanotube dissemination in the tissue is due to combination of their unique aspect ratio and nanometer scale diameter. Their excellent photostability enabled individual nanotubes to be imaged at video rate for extended times (tens of minutes) with neither evidence of photobleaching nor visible photodamage to the tissue observed.

Figures 5A and 5B are characteristic images of unf- and f-SWCNTs located at depths of more than 15 μm inside the live brain slices. For this experiment 845 nm excitation was adjusted *in situ* to 2,500 W/cm^2^ to excite unf-SWCNTs while only 100 W/cm^2^ was necessary to efficiently excite f-SWCNTs as displayed in Figure 5. More quantitatively, by comparing SWCNTs displaying similar signals (i.e. ∼500 camera counts), the corresponding signal to noise ratio (S/N, as average value evaluated for 10 individual nanotubes) were ∼ 7 for unf-SWCNTs excited at 845 nm and ∼ 11 for f-SWCNTs excited at 985 nm (Figure 5C & 5D). Note that f-SWCNTs excited at 845 nm with 1.3 kW/cm^2^ displayed a S/N of ∼ 9. Several factors contribute to this better performance for f-SWCNTs. First, light attenuation, scattering and background noise are minimized in the NIR-II region (1000-1700 nm), in which f-SWCNTs emit. More precisely the primary contributions to elevated background signals in NIR-I as compared to NIR-II are scattering of emitted photons which degrade image formations by ballistic photons, and autofluorescence from biological media. Moreover, the large absorbance cross-section of the E_11_ transition lowers the required excitation intensity, which consequently reduces light scattering, absorption, and tissue autofluorescence that results from excitation. Note that the blurry signals, particularly visible in Figure 5B, correspond to out-of-focus f-SWCNTs and not cellular tissue background as they can be resolved by changing the focus at the camera. Altogether, combination of more efficient excitation and improved imaging in the NIR-II region led to better S/N PL imaging ratio using less than 20 times the light excitation dose for imaging single f-SWCNTs which is an obvious advantage for non-invasive bioimaging.

**Figure 5:**
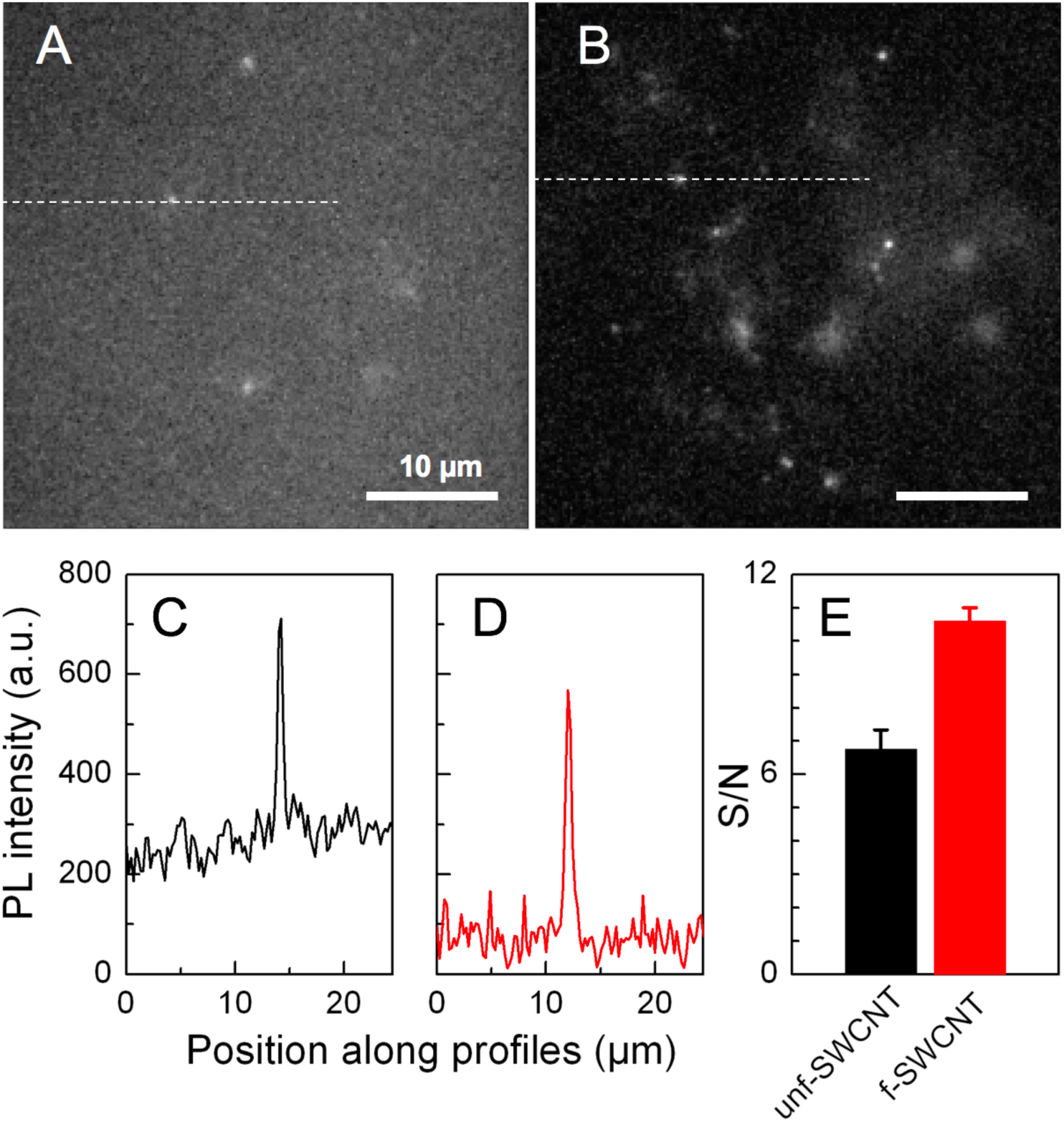
Live brain tissue imaging using SWCNTs. (A) unf-SWCNTs (845 nm excitation; 2.5 kW/cm^2^) and (B) f-SWCNTs (985 nm excitation; 100 W/cm^2^) are detected at depths >15 μm inside live brain slices. (C) and (D) Image section along the profiles (dashed lines) indicated in (A) and (B), respectively for unf-SWCNTs (red) and f-SWCNTs (black) display equivalent PL intensity at their respective emission wavelengths using different excitation intensities. (E) Signal-to-noise (mean ± stdev) values for unf- and f-SWCNT were each obtained from 10 individual nanotubes.

## Conclusion

Sensitive fluorescence microscopy deep within live biological samples using minimal light excitation dose for broad applications of high-resolution fluorescence approaches in biological science has been demonstrated using chemically modified SWCNTs. The synthesized f-SWCNTs are biocompatible and fluoresce brightly in the NIR-II due to the incorporated sp^3^ defects, making them suitable for single nanotube imaging in live brain tissues. We quantitatively demonstrated single nanotube imaging in the NIR-II region at high signal-to-noise ratios using unprecedentedly low excitation intensities (100 W/cm^2^) that are an order of magnitude lower than those previously reported for SWCNTs. This work paves the way toward the application of sp^3^ defect-tailored nanotubes as single-molecule probes ^[39]^ and chemical/molecular sensors ^[15,40]^ in live biological tissues.

## Acknowledgements

The University of Bordeaux and the Centre National de la Recherche Scientifique provided infrastructural support. This work was supported in part by grants from the Agence Nationale de la Recherche (ANR-14-OHRI-0001-01, ANR-15-CE16-0004-03), IdEx Bordeaux (ANR-10-IDEX-03-02), the France-BioImaging national infrastructure (ANR-10-INBS-04-01), and the Labex Brain Program (ANR-10-LABX-43) to L.C. and L.G., and by NSF (CHE-1507974) and NIH/NIGMS (R01GM114167) to Y.H.W. J.S.F. would like to thank the IINS Cell Culture Facility, particularly Emeline Verdier for the help with the organotypic slices preparation. Animal experiments were performed at the Animal Facilities of the University of Bordeaux, supported by the Région Nouvelle-Aquitaine.

